# Characterization of long non-coding RNAs during compatible and incompatible pollination in *Arabidopsis thaliana*

**DOI:** 10.64898/2026.04.29.721561

**Authors:** Nischay Patel, Nilesh D. Gawande, Subramanian Sankaranarayanan

## Abstract

Long non-coding RNAs (lncRNAs) have emerged as critical players in plant development and stress responses, yet their involvement in pollination responses is largely unknown. To address this gap, we identified and characterized lncRNAs and their cis-acting, trans-acting, and miRNA-mediated regulatory interactions during both compatible and incompatible pollination in *Arabidopsis thaliana*. Leveraging publicly available datasets, we analyzed expression profiles at 10 and 60 minutes post-pollination. We identified 1,073 novel and 3,422 annotated lncRNAs, with 1,002 novel and 985 annotated, respectively, showing detectable expression after filtering. Differential expression analysis identified 12 lncRNAs at 10 min and 32 lncRNAs at 60 min post-pollination. Further investigation revealed 9 cis-targets, 112 trans-targets, and 144 miRNA-mediated regulatory interactions, many of which were enriched in pathways related to stress, defense, and self-incompatibility. Notably, the regulatory landscape is more active at 60 minutes than at 10 minutes post-pollination. These findings provide a robust framework and resource to facilitate future functional studies of lncRNAs during pollination.

## 1. Introduction

Long non-coding RNAs (lncRNAs) are characterized by transcripts longer than 200 base pairs that do not encode proteins. Plant genomes consist of tens of thousands of lncRNAs that originate from coding regions, introns, or intergenic regions, and are usually transcribed by RNA polymerase II from both the sense and antisense strands. However, other polymerases, such as RNA polymerase IV and V, also produce lncRNAs in plants (Wierzbicki et al., 2021). LncRNAs play a key role in the regulation of molecular and biological processes, as well as cellular and developmental functions, through RNA-dependent mechanisms.

LncRNAs exhibit diverse regulatory functions, determined by their sequence composition, structure, and interaction partners (Wang and Chang, 2011; Wierzbicki et al., 2021). In plants, lncRNAs are involved in various cellular and molecular mechanisms, including chromatin modification, transcriptional regulation, RNA processing, mRNA stability, and translational control (Ariel et al., 2015; Quinn and Chang, 2016). LncRNAs exhibit distinct characteristics, such as sequence conservation, lower expression levels, and varied treatment-and tissue-specific expression patterns, compared to protein-coding genes. Beyond these, lncRNAs often play regulatory roles (Ulitsky 2016; Golicz et al., 2018).

In plants, lncRNAs are categorized based on their relative genomic location to protein-coding genes. Sense lncRNA overlaps with the exonic regions of protein-coding genes on same strand, while antisense lncRNAs are transcribed from the opposite strand of protein-coding genes. Intronic lncRNAs are transcribed from intronic regions of protein-coding genes, while intergenic lncRNAs are produced from regions between the annotated genes. LncRNAs can function either by acting near the genomic locus from which they are transcribed (cis) or by regulating distant targets (trans), including through miRNA-mediated regulatory interactions such as target mimicry, where lncRNAs bind miRNA and block their interaction with target mRNA (Franco-Zorrilla et al., 2007; Gil and Ulitsky, 2020).

In flowering plants, successful fertilization determines the future of seed production. Various signaling events happening during communication between compatible male reproductive tissue (pollen) and the female tissue (the pistil) have a critical role in reproduction. During pollination, a desiccated pollen lands on the stigma papillae cells of the pistil and shuts down the Reactive Oxygen species (ROS) signaling pathway at the stigmatic surface, resulting in further events such pollen hydration and germination, which further leads to the formation and elongation of the pollen tube through the stigmatic cell wall, enabling it to reach the ovary and release the male gametes to the ovule for fertilization (Jamshed et al., 2023). Plants distinguish between compatible and incompatible pollen, which helps them achieve reproductive success and maintain genetic diversity (Muñoz-Sanz et al., 2020).

In Brassicaceae species such as Arabidopsis and Brassica, compatible pollination involves several steps, including pollen hydration, germination, and pollen tube growth. In contrast, incompatible pollination leads to rapid pollen rejection at the stigma surface itself (Hiscock 2002; Chae and Lord 2011). Self-incompatibility responses are highly specific and are mediated by interaction between the stigma-expressed S-locus receptor kinase (SRK) and the pollen coat-derived S-locus cysteine-rich protein (SCR/SP11) in Brassicaceae, resulting in activation of signaling cascades that lead to pollen germination inhibition (Takayama et al., 2000; Yamamoto and Nishio, 2014; Bhalla et al., 2025a). The self pollen or a foreign pollen interaction with stigmatic papillae initiates a unique transcriptional response within early minutes of pollination, which eventually decides whether pollen is accepted or rejected (Sankaranarayanan et al., 2013; Zhang et al., 2017; Kodera et al., 2021). This selective process is biologically significant as it is a key determinant of fertilization efficiency and seed set (Ferrer et al., 2009). Therefore, the transcriptional reprogramming in pollen and stigma during compatible and incompatible pollination includes various cellular responses, signaling events, vesicle trafficking, cytoskeletal remodelling, and metabolic adjustments that determine pollen acceptance or rejection (Iwano et al., 2007; Kodera et al., 2021).

At the molecular level, transcriptome analyses in Arabidopsis and *Brassica napus* demonstrate that pollination responses are highly dynamic and time-dependent. Early transcriptional changes are detectable as soon as 10 minutes after pollination and diverge into distinct expression profiles by 60 minutes, distinguishing compatible from incompatible interactions (Sankaranarayanan et al., 2013; Zhang et al., 2017; Kodera et al., 2021). In the stigmas, during the compatible pollination, genes involved in MAPK signaling and plant-pathogen interaction pathways such as *Mitogen-Activated Protein Kinase 3* and *6* (*MKK3/6*) and *WRKY Transcription Factor 22* and *33* (*WRKY22/33*), are rapidly induced within 10 min, with particularly strong upregulation of *MPK3* and *WRKY33*. In contrast, incompatible pollination elicits only minimal expression of a few stigma-specific genes at 10 minutes; however, by 60 minutes, more than 100 genes are upregulated, indicating a delayed yet extensive transcriptional response characteristic of incompatibility (Kodera et al., 2021). The incompatibility-related genes upregulated at 60 min include receptor-like kinases *CRK41* and *CRK31*, and factors associated with endocytosis, secretion, and defense-related processes, such as *FLOT1*, *ROH1*, and *MLO12*. In pollen, compatible interactions are characterized by the upregulation of pollen-expressed and pollen tube associated genes, including *CHX21*, implicated in pollen tube guidance, as well as a receptor-like cytoplasmic kinase *(RLCK)* family gene at 60 min, both showing higher fold changes compared to incompatible interactions at the same time point (Kodera et al., 2021). These varied transcriptional changes across time points highlight the significance of time-dependent sample collection in capturing early versus late regulatory molecular signatures (Sankaranarayanan et al., 2013; Zhang et al., 2017).

In plant growth regulation, long non-coding RNAs (lncRNAs) play diverse functional roles, including antisense and splicing-associated regulation. Notable examples such as COOLAIR and ASCO contribute to the maintenance of shoot apical meristem integrity by modulating transcriptional programs, alternative splicing, and auxin-responsive gene expression (Marquardt et al., 2014; Rosa et al., 2016; Rigo et al., 2020). Another auxin-responsive lncRNA, APOLO, functions in roots to regulate lateral root and root hair development by translating auxin signals into chromatin-level modifications, thereby reprogramming transcriptional networks that control auxin distribution and root patterning (Ariel et al., 2014; Ariel et al., 2020). Additionally, the antisense lncRNA associated with DOG1 regulates seed dormancy through chromatin-mediated transcriptional control (Fedak et al., 2016). Beyond their established roles in plant development and hormone signaling, these findings suggest that lncRNAs may also contribute to reproductive processes, particularly through transcriptional reprogramming during pollen–pistil interactions.

In plant reproduction, functional evidence supporting the role of lncRNAs in pollen fertility has been demonstrated in rice, where disruption of the lncRNA LDMAR leads to photoperiod-sensitive male sterility, directly linking lncRNA regulation to reproductive success (Ding et al., 2012). High-throughput sequencing analyses of anthers across various developmental stages in *A. thaliana* identified 1,283 lncRNAs, with predicted functions including roles as miRNA precursors, endogenous target mimics, or natural antisense transcripts of primary miRNA (Zhou et al., 2025). Similarly, in *Brassica rapa*, 12,051 lncRNAs were profiled across five pollen stages, including pollen mother cell, tetrad, uninucleate pollen, binucleate pollen, and mature pollen (Huang et al., 2018). Environmental cues further influence lncRNA activity during reproductive development; for example, heat stress during pollen development in wheat induces 5,482 lncRNAs associated with predicted cis-and trans-target genes involved in heat response, protein folding, abiotic stress signaling, and jasmonic acid biosynthesis pathways (Babaei et al., 2024).

Despite the growing body of transcriptomic data, most studies have focused primarily on protein-coding genes, leaving the roles of non-coding regulators such as lncRNAs during pollination relatively underexplored. Although lncRNAs are increasingly recognized as key regulators of gene expression, their specific contributions to pollination and reproductive responses remain poorly understood.

In this study, we identified and characterized lncRNAs using transcriptomic datasets from pollinated Arabidopsis stigmas under compatible and incompatible pollination conditions at 10 and 60 min, respectively (Kodera et al., 2021). The identified lncRNAs were classified based on their genomic context, specific expression patterns, and differential expression at 10 min and 60 min after pollination. Furthermore, we predicted cis-and trans-regulatory interactions between lncRNAs and protein-coding genes, constructed lncRNA–miRNA–mRNA regulatory networks, and performed functional enrichment analysis to infer the potential biological roles. Together, these findings provide new insights into the spatial and temporal dynamics of lncRNA expression in stigma and pollen tissues during compatible and incompatible pollination. This study also highlights the interactions between lncRNAs and other regulatory miRNAs, elucidating the complex regulatory networks that underpin pollination-associated transcriptional reprogramming.

## 2. Materials and methods

### 2.1. Data retrieval and Quality Control

The raw RNA-seq data for compatible and incompatible pollination at two time points (10 min and 60 min post-pollination were retrieved from a publicly available dataset (Accession no: SRP154565, Kodera et al., 2021) in the Sequence Read Archive (SRA) at the National Center for Biotechnology Information (NCBI) database. The dataset included the transcriptomes of pollinated stigmatic tissues from compatible pollination (Col-0 /*SRK14* x C24) and incompatible pollination (Col-0 /SRK14 x C24/SCR14) (Kodera et al., 2021) at 10 min (t = 10) and 60 min (t = 60) post-pollination (**Supplementary Table S1**). The reference genome sequence and gene annotation files for *A. thaliana* were obtained from the Ensembl Plants database (https://plants.ensembl.org/index.html). The details for the samples and reference annotation file sources are provided in **Supplementary Tables S1 and S2**.

RNA-seq data quality and replicate consistency were assessed using FastQC (Andrews, 2010) and principal component analysis (PCA) of variance-stabilized expression values. This analysis was further complemented by a Euclidean distance-based metric calculated on VST-transformed data to assess the relatedness among biological replicates **(Supplementary Table S3 and Supplementary Figure S1)**. The final dataset included three independent biological replicates for pollinated stigmas under compatible and incompatible pollination, collected at 10 and 60 min post-pollination.

### 2.2. Transcriptome reconstruction and lncRNA identification pipeline

The adapter sequences and low-quality bases in the raw reads were trimmed using Trimmomatic (Bolger et al., 2014). The library strandedness for the RNA-seq samples was determined using an online tool (https://github.com/signalbash/how_are_we_stranded_here) (Signal and Kahlke 2022). High-quality reads were aligned to the *A. thaliana* reference genome from TAIR10 using HISAT2 (Kim et al., 2015). For each sample, per-sample transcript assemblies were generated using StringTie (Pertea et al., 2015; Kovaka et al., 2019) in reference guided mode. Six independent assemblies per time point were merged into a unified transcriptome annotation with StringTie-merge, and the merged transcriptome was compared to the TAIR10 reference annotation using GffCompare (Pertea and Pertea, 2020) to classify transcript structures. Reference transcripts were linked to TAIR10 identifiers, while novel transcripts were assigned locus-level (XLOC) and transcript-level (TLOC) identifiers. Further, the merged annotation in GTF format was converted to BED12 format, and transcript sequences were extracted from the TAIR10 genome using the BEDTools getfasta function (Quinlan and Hall, 2010). Transcripts shorter than 200 nucleotides were excluded from further analysis. The protein-coding genes, small non-coding RNAs, and non-translating CDS were excluded from further analysis. Transcripts annotated as long non-coding RNAs (lncRNAs) in TAIR10 were retained as annotated lncRNAs, while the transcripts lacking lncRNA annotation in TAIR10 (transcripts annotated as non-coding RNAs) and previously unannotated transcripts were collectively treated as putative novel lncRNAs and were subjected to a further multi-step filtering pipeline for novel lncRNA identification. This approach identifies previously unannotated lncRNA candidates while avoiding reclassification of well-characterized protein-coding transcripts and small non-coding RNAs.

#### 2.2.1 Coding potential assessment

The coding potential of unannotated transcripts was determined using three independent tools that included CPC2, CPAT, and LncFinder. The cutoff considered for these tools included CPC2 (coding probability<0.5) (https://github.com/gao-lab/CPC2_standalone) (Kang et al., 2017), CPAT (coding probability<0.46) (https://github.com/liguowang/cpat) (Wang et al., 2013), and LncFinder (https://cran.r-project.org/web/packages/LncFinder/index.html) (coding potential<0.5) (Han et al., 2019). Plant-specific pre-trained models and corresponding cutoff values as defined in the Plant-LncPipe framework were used for CPAT and Plant LncFinder (https://github.com/xuechantian/Plant-LncRNA-pipline) (Tian et al., 2024). Transcripts that passed all three tools criteria were retained as novel candidate lncRNAs.

#### 2.2.2 Protein domain filtering

To further refine the selection of potential lncRNAs, novel candidate lncRNAs were screened against the Pfam database (Mistry et al., 2021) using PfamScan (HMMER) https://github.com/aziele/pfam_scan). Transcripts that contained Pfam domain hits with domain E-values ≤ 1E^-3^ were considered protein-coding and excluded from subsequent analyses.

#### 2.2.3 Construction of the final lncRNA set

The resulting high-confidence novel lncRNAs were integrated with previously annotated lncRNAs to create a final lncRNA set. From transcript-level annotations, lncRNA loci were defined by grouping lncRNA transcripts based on their genomic coordinates; any locus that generated at least one lncRNA transcript was classified as a lncRNA locus.

### 2.3. Classification of lncRNAs based on their genomic positions

The novel lncRNAs were subjected to the FEELnc (Flexible Extraction of Long non-coding RNAs) classifier (https://github.com/tderrien/FEELnc) (Wucher et al., 2017), which assigns lncRNAs to genomic categories based on their positional relationship to annotated protein-coding transcripts. Each lncRNA was first classified as either genic (overlapped an annotated gene) or intergenic (located between annotated genes). Genic lncRNAs were further subclassified based on their overlap with coding genes (exonic, intronic, overlapping, containing, or nested), whereas intergenic lncRNAs were categorized based on their relative orientation and genomic position relative to neighboring genes (divergent, convergent, or same-strand, and upstream or downstream).

### 2.4. Transcript expression quantification and classification

Transcript-level expression quantification was performed using Kallisto (Bray et al., 2016) using the merged transcriptome as the reference index. Expression levels for samples were calculated as transcripts per million (TPM). Transcripts and coding genes with low expression were filtered using mean–variance trend analysis. The inspection of the mean–variance relationship using LOWESS smoothing indicated increased variability at TPM ≤ 0.05 **(Supplementary Figure. S1)**. Considering the lower expression levels of lncRNAs compared to messenger RNAs (mRNAs) (Grammatikakis and Lal 2022), a permissive cutoff for expression of lncRNA transcripts was set at TPM > 0.05.

To ensure accurate expression analysis and minimize artifacts from low or inconsistent transcript detection, an expression-based filtering step was applied before differential expression analysis. An lncRNA was considered expressed at time points (t = 10 or t = 60 min) if at least one transcript had TPM > 0.05 in at least two of three biological replicates in either the compatible or incompatible condition. Among the expressed lncRNAs, condition-specific expression was defined using a stricter criterion: lncRNA transcripts were classified as condition-specific if they had TPM > 0.1 in at least two out of three replicates in one condition and no detectable expression (TPM = 0) in all three replicates of the other condition at the same time. The expressed lncRNA genes and their corresponding transcripts were retained for differential expression analysis.

### 2.5. Differential expression analysis of lncRNAs and mRNAs

Gene level counts and transcripts per million (TPM) values were generated from transcript-level estimates using the tximport R package with the lengthScaledTPM method. Differential expression analysis of expressed lncRNA genes and protein-coding mRNAs was conducted using DESeq2 (Love et al., 2014). Fold change was calculated as, Fold Change = TPM (Incompatible Sample) / TPM (Compatible Sample). Statistical significance was assessed with Wald tests, and p-values were adjusted for multiple testing using the Benjamini-Hochberg false discovery rate (FDR) (Haynes 2013). Differential expression was considered significant for adjusted p-values < 0.1 and absolute log_2_ fold changes ≥ 1. Subsequent functional analysis and biological interpretation were performed at the gene (locus) level.

### 2.6. Identification of cis and trans targets of lncRNAs

Cis target genes of lncRNAs were identified based on genomic proximity and expression correlation, focusing only on differentially expressed (DE) lncRNAs. DE protein-coding mRNAs were treated as potential DE lncRNA targets. For each lncRNA, the ten nearest protein-coding genes located within 100 kb upstream and downstream of the lncRNA locus were identified using BEDTools (Quinlan and Hall 2010). Overlapping protein-coding genes were also included. Genes that did not fall within a 100 kb were considered as potential trans targets. To refine the lncRNA-mRNA associations, Pearson correlation coefficients and their p-values were calculated for each candidate lncRNA-mRNA pair using the R stats package (cor.test function). LncRNA–mRNA pairs that met the criteria of |r| > 0.81 and adjusted p-value ≤ 0.05 were considered as putative cis interactions of lncRNAs.

Protein-coding genes located more than 100 kb from the lncRNA locus were considered as potential trans targets. Trans target identification was performed on protein-coding genes using the same correlation framework and multiple-testing correction was applied across the tested trans-acting lncRNA–mRNA pairs using the Benjamini–Hochberg false discovery rate (FDR) method (Haynes 2013). LncRNA–mRNA (|r| > 0.81 and p-value ≤ 0.05) were retained as putative trans-acting associations. Trans interactions were validated using LncTar (Li et al., 2015), which estimates the normalized binding free energy (ndG) between co-expressed lncRNAs and their paired protein-coding genes. Given the non-canonical nature of lncRNA–mRNA interactions in plants, only lncRNA–mRNA interaction pairs with ndG ≤ −0.08 were retained as energetically plausible interactions, as reported previously (Li et al., 2015).

### 2.7. Prediction of lncRNA-miRNA-mRNA regulatory network

To infer the functions of DE lncRNAs, a lncRNA–miRNA–mRNA regulatory network was constructed based on predicted RNA–RNA interactions. Mature miRNA sequences were obtained from the miRBase database (Kozomara et al., 2019). The psRNATarget tool (Dai et al., 2018) was used to predict interactions between DE lncRNAs and miRNAs, as well as between miRNAs and DE protein-coding mRNAs, applying an expectation cut-off of ≤ 3.5 for miRNA–mRNA and ≤ 4.5 for lncRNA–miRNA interactions. The resulting interaction pairs were used to construct the regulatory network and were visualized using Cytoscape (Shannon et al., 2003).

### 2.8. Functional prediction and enrichment analysis of lncRNAs

Functional prediction of lncRNAs was based on their associated protein-coding target genes. The DE target mRNAs identified from cis-and trans-acting lncRNA associations and the lncRNA–miRNA–mRNA regulatory network were used for enrichment analysis. The Arabidopsis TAIR10 genome annotation was used to construct the background gene set. Gene Ontology (GO) term analysis was conducted using the clusterProfiler package in R (Yu et al., 2012), with statistical significance assessed at an adjusted p-value (Padj ≤ 0.05). Additionally, the functional descriptions of the target genes were annotated using publicly available TAIR10 resources, and a heatmap was generated in R using the Complex Heatmap package (Gu et al., 2016).

### 2.9. Computational environment and data processing

Data processing, statistical analyses, and result integration were performed in the R environment using RStudio. RNA-seq analysis, including read processing, was conducted on the Galaxy server (Blankenberg et al., 2014), while downstream handling, filtering, statistical testing, and visualization were done locally in R. Plots were generated with common libraries like ggplots and VennDiagram (Chen and Boutros, 2011), and schematic illustrations were created using BioRender. Custom Bash scripts were used for specific data preprocessing tasks via the command line. An overview of the RNA-seq workflow is in **Supplementary Table S4**, and the software tools used, along with their versions, settings, and computational platforms, are detailed in **Supplementary Table S5**.

## 3. Results

### 3.1. Sense genic lncRNA transcripts are predominant among expressed novel lncRNAs

LncRNAs associated with compatible and incompatible pollination responses were determined from RNA-seq data of pollinated *A. thaliana* stigmatic tissues using the workflow illustrated in **Figure 1**. The dataset comprised a total of 12 RNA-seq samples of pollinated stigmatic tissues from compatible and incompatible pollination at early (t = 10 min) and late (t = 60 min) post-pollination stages, with three biological replicates per condition and time point. Only paired-end reads that passed the filtering criteria were retained for further analysis. The proportion of discarded read pairs ranged from 1.72% to 2.46% across the samples (**Supplementary Table S6**). The alignment of high-quality reads to the A. thalian TAIR10 reference genome using HISAT2 displayed alignment rates of more than 97% across the samples (**Supplementary Table S7**), indicating high-quality sequencing and mapping.

**Figure 1.**
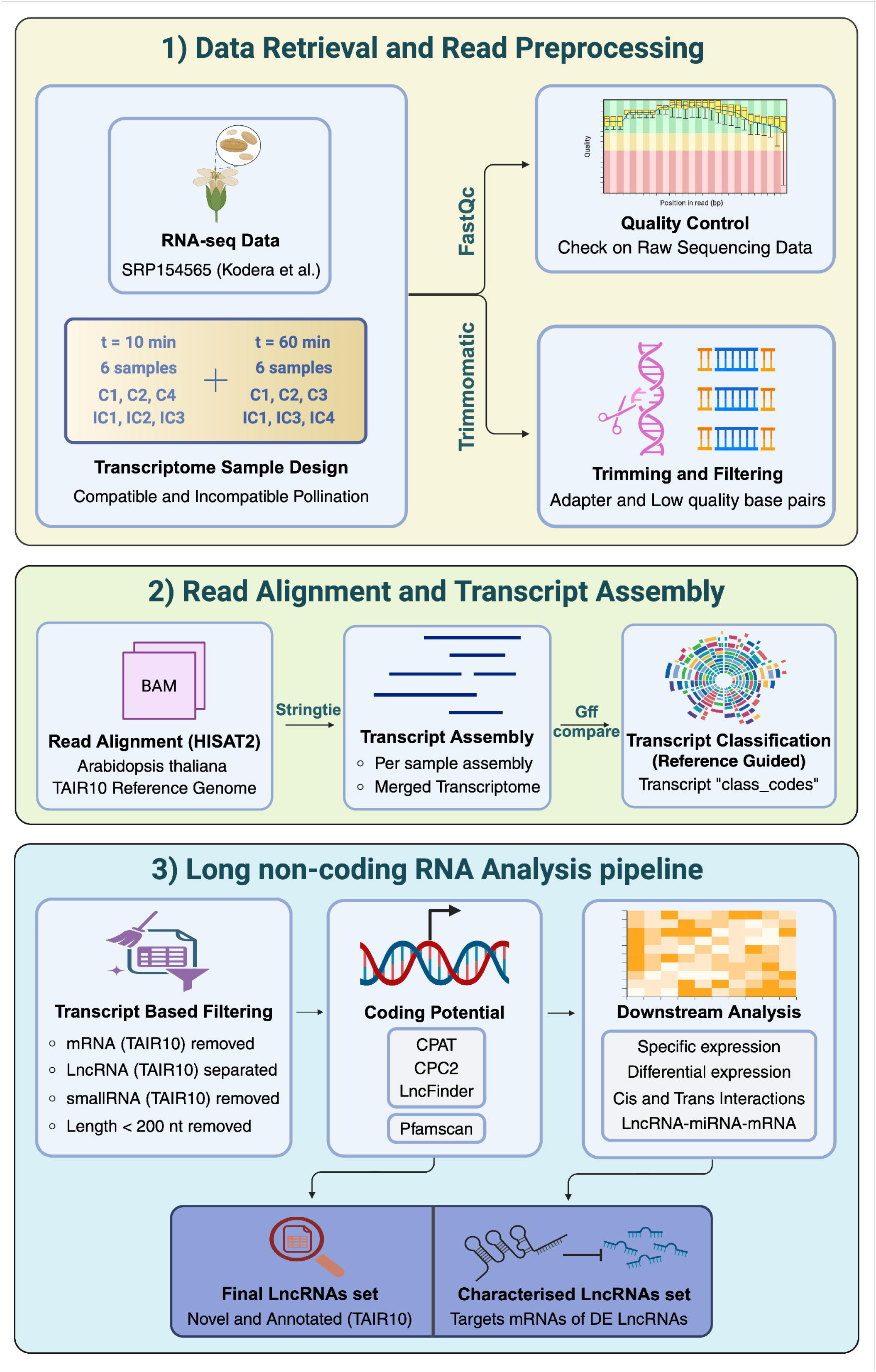
**Overview of the RNA-seq data workflow for processing, identifying, characterizing, and analyzing long non-coding RNAs.**

The merged transcriptome consisted of 63,455 transcripts corresponding to 37,792 genomic loci **(Figure 2a)**. After filtering for transcripts longer than 200 bp, 61,836 transcripts (36,192 loci) were retained. The removal of known protein-coding transcripts and annotated non-lncRNA biotypes (tRNA, rRNA, snRNA, snoRNA, and miRNA) resulted in 9,888 candidate transcripts that were subjected to coding-potential assessment. Coding potential assessment using three independent tools, CPAT, CPC2, and LncFinder, effectively identified 1,533 novel lncRNA transcripts associated with 1,073 lncRNA loci **(Figure 2b).** None of these transcripts displayed specific functional or structural protein domains using the Pfam database **(Supplementary Table S8)**. In total, 1,381 novel lncRNA transcripts derived from 1,002 expressed lncRNA loci met the TPM and replicate consistency criteria. In addition, 1,260 annotated lncRNA transcripts from 985 expressed lncRNA loci were identified from TAIR10, resulting in a comprehensive set of expressed novel and annotated lncRNAs **(Figure 2c and Supplementary Table S9)**.

**Figure 2.**
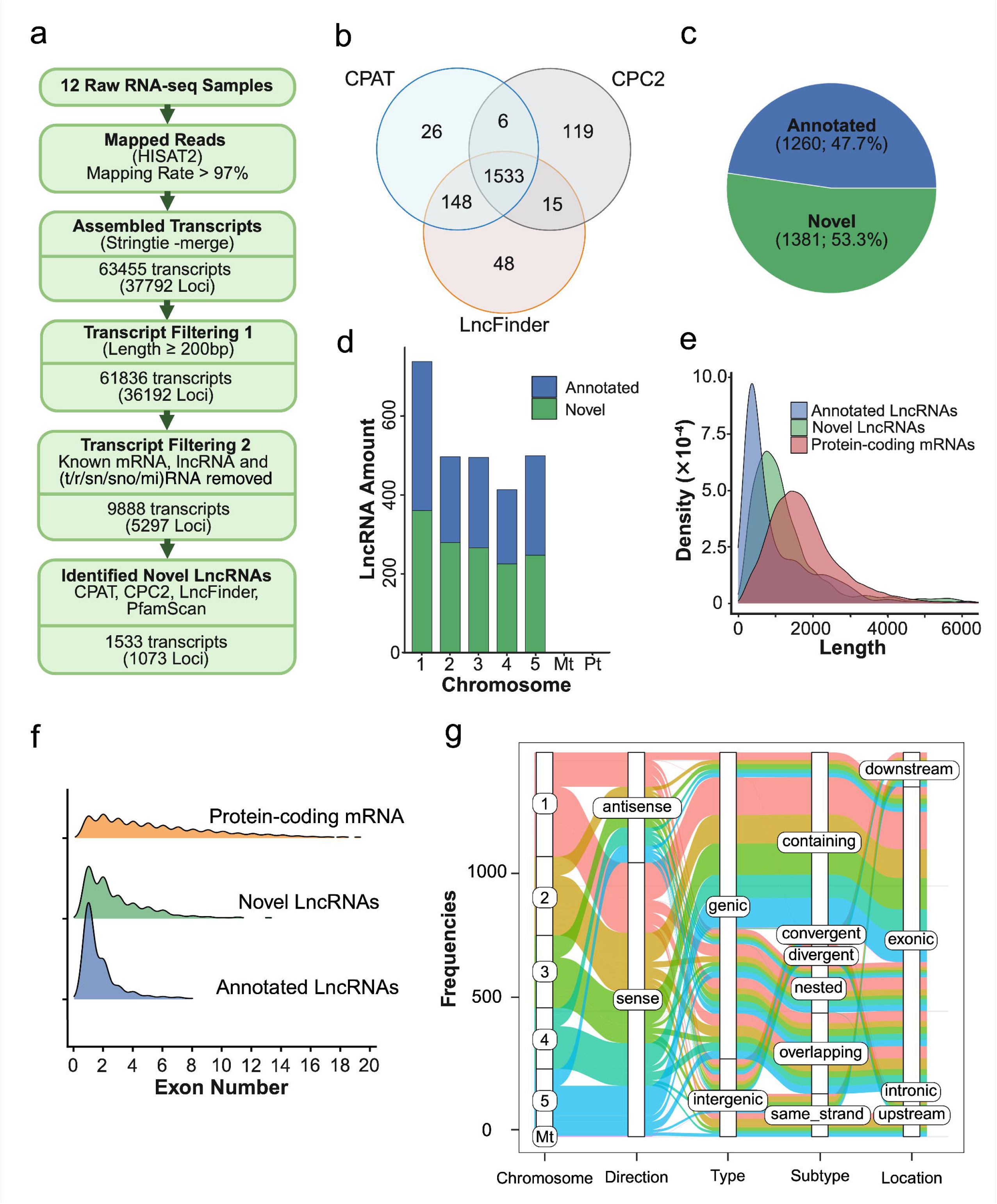
Characteristic features of annotated and novel long non-coding RNAs identified in pollinated stigmatic tissues of compatible and incompatible pollination in *Arabidopsis thaliana* at early (t = 10 min) and late (t = 60 min) post-pollination. The figure denote, (a) Schematic overview of the bioinformatic workflow used for RNA-seq processing, transcript assembly, sequential filtering, and identification of high-confidence novel lncRNAs (b) Venn diagram showing the overlap of putative non-coding transcripts predicted by CPAT, CPC2, and LncFinder (c) Proportion and absolute numbers of annotated and novel lncRNA-producing loci among expressed lncRNAs (d) Chromosomal distribution of annotated and novel lncRNA-producing loci (e) Length distribution of expressed annotated lncRNAs, expressed novel lncRNAs, and expressed protein-coding mRNAs (f) Exon number distribution of expressed novel lncRNAs, expressed annotated lncRNAs, and expressed protein-coding mRNAs (g) Genomic classification of expressed novel lncRNAs based on positional relationship to neighboring protein-coding genes

The expressed lncRNAs loci were unevenly distributed on chromosomes, with the highest number associated with chromosome 1 **(Figure 2d)**. The novel lncRNAs were generally shorter than annotated lncRNAs and protein-coding mRNAs, with mean transcript lengths of 1,186 bp for lncRNAs and 1,830 bp for protein-coding transcripts **(Figure 2e)**. Consistent with known features, lncRNAs contained fewer exons than protein-coding genes, while novel lncRNA transcripts exhibited a higher exon count compared with annotated lncRNA transcripts **(Figure 2f)**. Detailed information on chromosomal distribution, transcript length, and exon number for both identified and expressed annotated and novel lncRNAs is provided in **Supplementary Tables S8 and S9**.

Genomic classification of the 1,381 expressed novel lncRNA loci revealed that 70% (972/1,381) of transcripts were sense genic lncRNAs, while 30% (409/1,381) were antisense lncRNAs. Among the antisense lncRNAs, 19.8% (274/1,381) were classified as exonic. Intergenic lncRNAs accounted for 21.2% (293/1,381), and only 0.6% (8/1,381) lncRNAs were intronic. Furthermore, 21% of transcripts (290/1,381) overlapped with protein-coding genes. Together, these results suggest that sense-genic lncRNAs were abundantly expressed among the novel lncRNAs **(Figure 2g)**. Detailed FEELnc-based genomic classification of novel lncRNAs is provided in **Supplementary Table S10**.

### 3.2 Late post-pollination is characterized by an increased number of DE lncRNAs

LncRNAs exhibited lower expression levels than protein-coding genes, as reflected by the distribution of log[(TPM) values **(Figure 3a)** (Grammatikakis and Lal 2022). Determination of the mean–variance relationship of transcript expression further revealed increased variability at low expression levels (TPM < 0.05) **(Supplementary Figure S2)**. Based on stringent criteria for TPM thresholds and replicate consistency, a total of 1,821 and 1,830 lncRNA loci were identified as expressed at t = 10 min and t = 60 min, respectively. Among these, 1,664 loci were expressed at both time points, whereas 157 loci were uniquely expressed at t = 10 min and 166 loci were specific to t = 60 min, respectively **(Figure 3b).** Thus, the expressed number of lncRNA loci was comparable between early and late post-pollination stages **(Supplementary Table S9)**.

**Figure 3.**
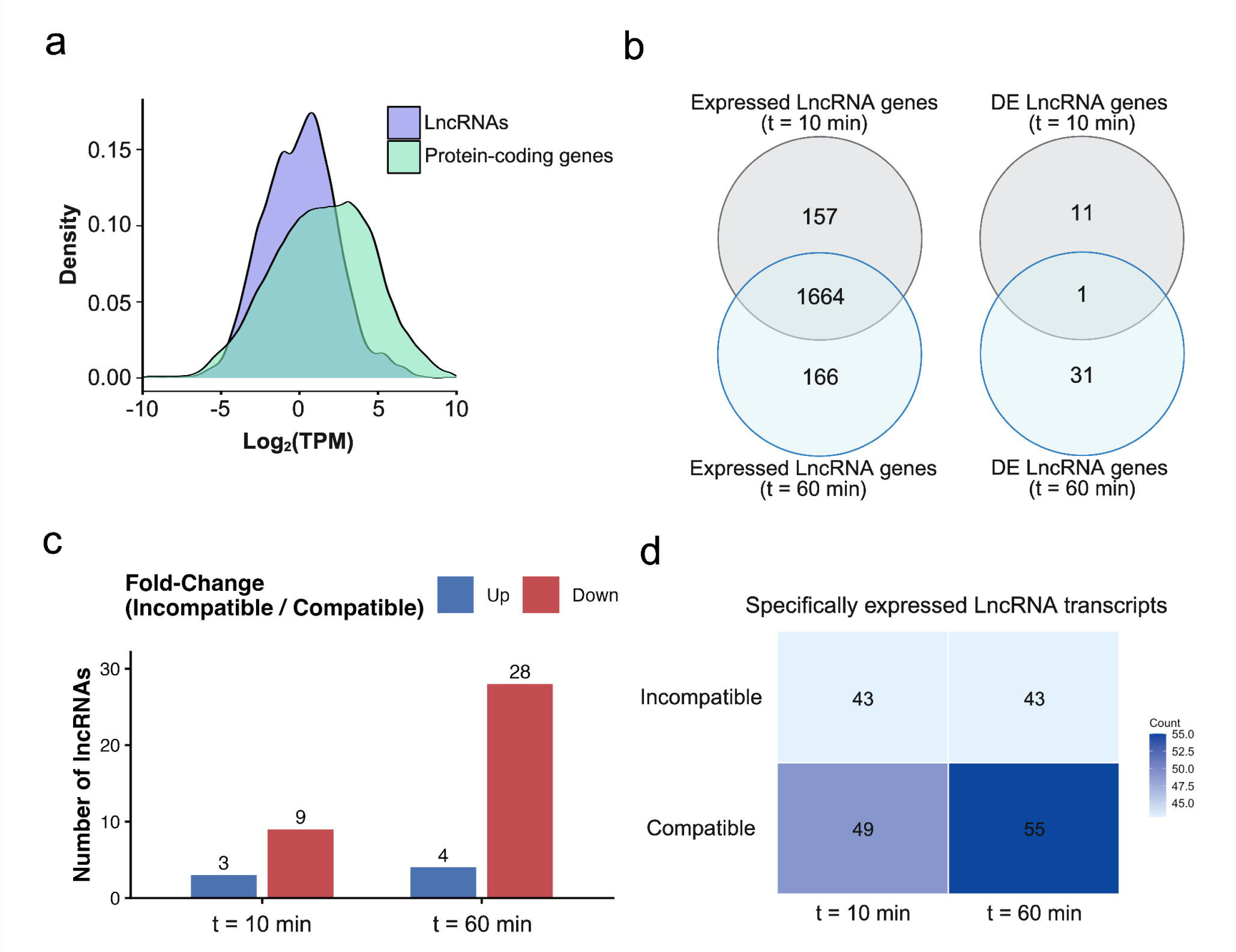
Expression classification and temporal expression pattern of lncRNAs in pollinated stigmatic tissues of compatible and incompatible pollination in *Arabidopsis thaliana* at early (t = 10 min) and late (t = 60 min) post-pollination. The figure displays (a) Density distribution of transcript expression levels showing log[-transformed TPM values for lncRNAs and protein-coding genes across all samples. LncRNAs exhibit lower overall expression levels than protein-coding genes. (b) Venn diagrams depicting the number of expressed lncRNA genes (left) and differentially expressed (DE) lncRNA genes (right) shared between t = 10 min and t = 60 min, as well as those specific to each time point. Differential expression was assessed as incompatible vs compatible pollination conditions. (c) Number of upregulated and downregulated DE lncRNAs at t = 10 min and t = 60 min, based on fold change. (d) Heatmap summarizing specifically expressed lncRNA transcripts, defined as transcripts expressed in one pollination condition (compatible or incompatible) and absent in the other at the same time point.

DE analysis of incompatible and compatible pollination (p-value < 0.1 and an absolute log_2_ FC Incompatible pollination/Compatible pollination ≥ 1) identified 45 DE lncRNA loci across both time points, out of which 37 loci met a stringent significance threshold (adjusted p < 0.05). The remaining loci were retained at a relaxed threshold (adjusted p < 0.1). Only a single DE lncRNA locus was shared between t = 10 min and t = 60 min, whereas 11 loci were specific to t = 10 min and 31 loci were specific to t = 60 min **(Figure 3b)**. Thus, despite broadly similar numbers of expressed lncRNA loci at both time points, a substantially larger number of DE lncRNAs was detected at 60 min, highlighting a temporal shift in lncRNA regulation from t = 10 min to t = 60 min post-pollination stage. The majority of DE lncRNAs were downregulated under incompatible pollination conditions. At t = 10 min, 9 out of 12 DE lncRNAs were downregulated, while at t = 60 min, 28 out of 32 DE lncRNAs showed downregulation. Overall, 37 of the 45 DE lncRNAs were downregulated under incompatible pollination **(Figure 3c)**. The most significant downregulation of DE lncRNAs was found at 10 min (log[fold change = −7.8), and a comparatively less downregulation at 60 min (−3.7). In contrast, mRNA showed a log[fold change of −8.5 at 10 min but strong upregulation of 9.8-fold at 60 min (9.8). The corresponding DE lncRNAs and protein-coding mRNAs are provided in **Supplementary Tables S11 and S12**, while DESeq2-normalized expression values are given in **Supplementary Tables S13A and S13B**.

Finally, lncRNA transcripts showing condition-specific expression were defined as those expressed in only one pollination condition (compatible or incompatible) at a given time point (t =10 min or t = 60 min). A higher number of specifically expressed lncRNA transcripts was observed under compatible pollination conditions at both t = 10 min and t = 60 min **(Figure 3d**, **Supplementary Tables S14A and S14B**).

### 3.3. Cis-regulatory associations of lncRNAs are enhanced in the late post-pollination stage

Cis interactions have been widely reported in plants, where lncRNAs modulate the expression of neighbouring genes through local transcriptional or chromatin-based mechanisms, enabling rapid and specific transcriptional responses during developmental and stress-related processes (Chekanova 2015; Lucero et al., 2021).

A total of 15 potential cis-acting lncRNA–mRNA pairs were analyzed across the two post-pollination time points. Among these, 10 significant cis interactions were identified, involving seven DE lncRNAs and nine DE mRNAs. A higher number of cis interactions was observed at the 60 min. The identified cis interactions exhibited strong expression correlations (|r| = 0.883-0.997). Of the ten significant cis pairs, eight interactions showed positive correlations, while two interactions showed negative correlations. Notably, six out of ten cis interactions exhibited very strong correlations (|r| > 0.95). Consistent with the direction of correlation, eight cis pairs displayed concordant regulation (both lncRNA and mRNA either upregulated or downregulated), whereas two pairs exhibited discordant regulation, characterized by opposite expression trends between the lncRNA and its neighbouring mRNA.

Cis-associated target mRNAs included genes such as *PCR2* (*AT1G14870*), *MAP3K13* (*AT1G07150*), and *DREB26* (*AT1G21910*), along with genes encoding pathogenesis-related thaumatin-like proteins, thionin-like peptides, chalcone–flavanone isomerase family proteins, phenolic glucoside malonyl transferase (PMAT1), SKU5-like proteins, and an ATPase F subunit. Among these, *PCR2* and a *thionin-like gene* were identified as overlapping targets with their associated lncRNAs, whereas the remaining cis interactions involved neighbouring but non-overlapping genes. The significant cis interactions, together with their genomic distances, correlation coefficients, statistical significance values, and functional annotations, are summarized in **Table 1** and **Supplementary Table 15**.

**Table 1.** Significant cis-acting lncRNA–mRNA interaction pairs identified at early (t = 10 min) and late (t = 60 min) pollination time points.

| Time Stage | lncRNA locus (IC/C) | mRNA Gene (IC/C) | Pearson Correlation | Distance | Target Description |
| --- | --- | --- | --- | --- | --- |
| t10 | XLOC_004982 (Down) | AT1G14870 (Down) | 0.973277004** | 0 | PCR2 encodes a membrane protein involved in zinc transport and detoxification. |
| t10 | XLOC_029258 (Down) | AT5G40020 (Up) | -0.883204065* | 15611 | Pathogenesis-related thaumatin superfamily protein;(source:Araport11) |
| t60 | AT1G07128 (Down) | AT1G07150 (Up) | -0.897819837* | -5545 | Member of MEKK subfamily. Involved in wound induced signaling where it interacts with At5g40440; and activates At1g59580. |
| t60 | AT3G51238 (Down) | AT3G51230 (Down) | 0.97631868** | 758 | chalcone-flavanone isomerase family protein;(source:Araport11) |
| t60 | XLOC_001211 (Down) | AT1G21860 (Down) | 0.95287363** | -1287 | SKU5 similar 7;(source:Araport11) |
|  |  | AT1G21866 (Down) | 0.905955412* | 0 | Thionin-like gene. |
|  |  | AT1G21910 (Down) | 0.946807793** | 18397 | encodes a member of the DREB subfamily A-5 of ERF/AP2 transcription factor family. The protein contains one AP2 domain. There are 15 members in this subfamily including RAP2.1; RAP2.9 and RAP2.10. |
| t60 | XLOC_004982 (Down) | AT1G14870 (Down) | 0.961194436** | 0 | PCR2 encodes a membrane protein involved in zinc transport and detoxification. |
| t60 | XLOC_029189 (Up) | AT5G39050 (Up) | 0.997231162** | -2017 | Encodes a malonyl transferase that may play a role in phenolic xenobiotic detoxification. |
| t60 | XLOC_031012 (Down) | ATCG00130 (Down) | 0.955603619** | 2064 | ATPase F subunit. |

**Note:** The table presents differentially expressed lncRNAs and their associated cis-target protein-coding genes, including the direction of expression change (up/down), Pearson correlation coefficients, genomic distances between lncRNA and target gene loci, and functional annotations of the target genes. Functional annotations were obtained from Araport11. Asterisks in the “Pearson correlation” column indicate significance levels (\*P < 0.05; \*\*P < 0.01). A comprehensive list of significant cis interactions is provided in **Supplementary Table 15**.

### 3.4. Trans-regulatory associations of lncRNAs are enhanced during the late post-pollination stage

The trans interaction landscape of DE lncRNAs was assessed using the distribution of ndG values for predicted lncRNA-mRNA pairs using a density plot **(Figure 4a)**. The trans interactions displayed a notable skew toward the right side of the chosen cutoff, indicating a high density of energetically permissible but relatively weaker interactions observed at ndG values 0.08 (ndG ≥-0.08). A significant decline in interaction density was observed around the ndG =-0.08 threshold. Most of the predicted interactions fell within the range (-0.08, - 0.1), and the number of interactions declined as the predicted interaction strength increased. A total of 118 significant trans-acting lncRNA-mRNA associations, consisting of 17 DE lncRNAs and 112 protein-coding genes, were found using combined expression correlation (|r| ≥ 0.81) and structural filtering (ndG ≤-0.08). Of these, 115 interactions were specific to the t =60 min, whereas only three interactions were detected at t = 10 min, indicating a strong enrichment of trans regulatory interactions at the later stage of pollination **(Supplementary Table 16)**.

**Figure 4.**
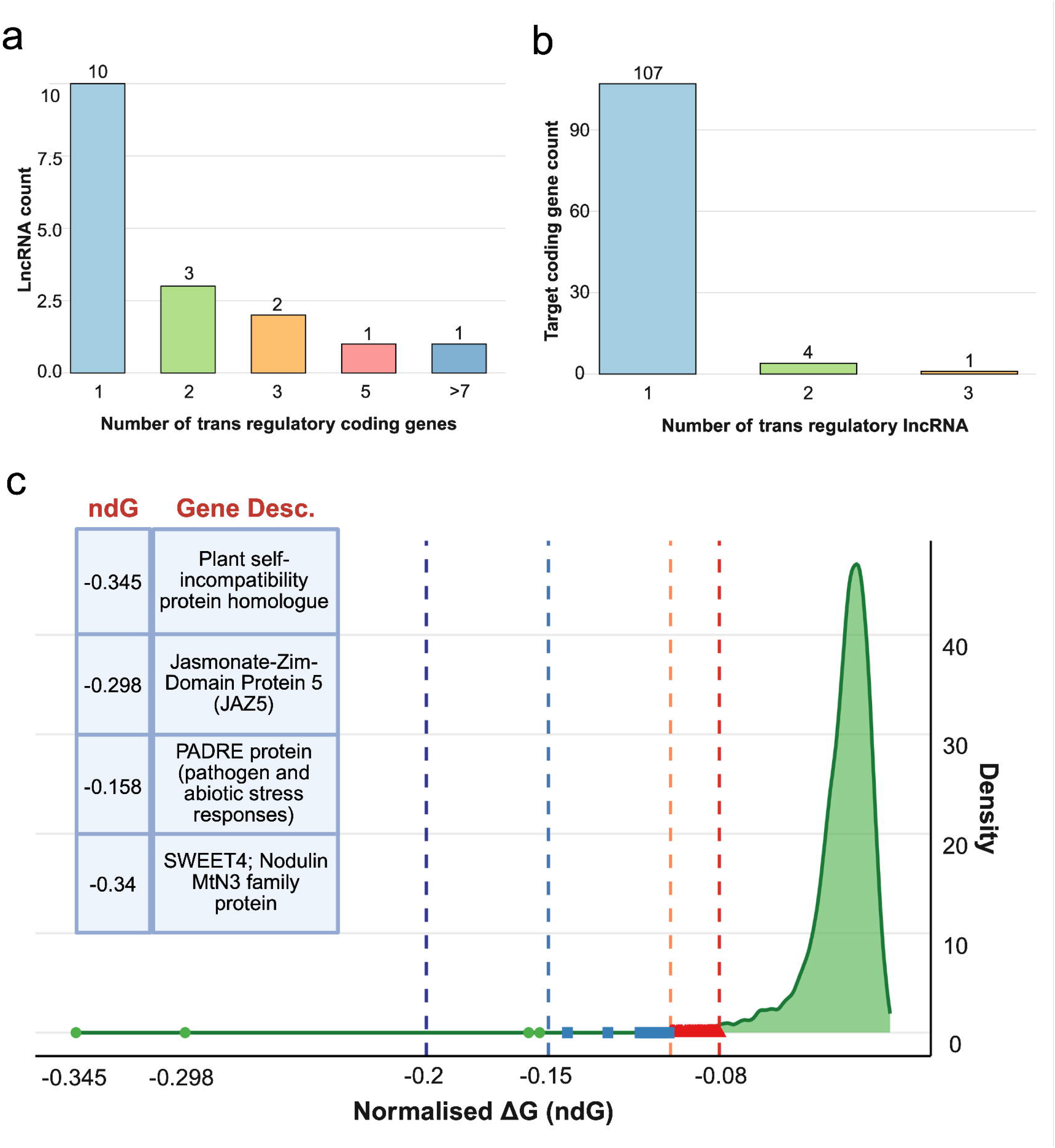
LncRNA–mRNA trans interaction landscape. The figure shows (a) Distribution of lncRNAs based on the number of predicted trans-regulatory protein-coding target genes. (b) Distribution of protein-coding genes according to the number of trans-regulatory lncRNAs. (c) Density distribution of normalized binding free energy (ndG) values for predicted lncRNA-mRNA trans interactions after filtering. The red dashed line indicates the ndG cutoff (≤-0.08). Interactions are categorized by ndG ranges: red triangles (-0.08 to - 0.10), blue squares (-0.10 to-0.15), and green circles (≤-0.15). The four targets with the most negative ndG values are highlighted.

To further investigate the regulatory network architecture of lncRNA-mediated trans regulation, the distribution of predicted trans target genes per lncRNA was analyzed **(Figure 4a)**. The majority of lncRNAs were associated with a single trans target, indicating that most lncRNAs exhibit limited trans-regulatory potential. Fewer lncRNAs targeted two or three genes, and only a very small subset exhibited higher target counts. Interestingly, four lncRNAs displayed the majority of predicted trans interactions, where the annotated lncRNA AT3G04795 was associated with 91 predicted protein-coding trans targets, while XLOC_017394, XLOC_001211, and XLOC_020056 were linked to five, three, and three targets, respectively. These four multi-target lncRNAs accounted for approximately 86% of all predicted trans interactions, suggesting that trans regulatory interactions were dominated by a few highly connected lncRNAs. The predicted targets for AT3G04795 included genes annotated as members of the plant self-incompatibility protein S1 family, the PADRE protein family, and the plant thionin protein family (Araport11). Of the 112 unique protein-coding genes identified as trans targets, 107 were associated with only a single lncRNA, indicating that most coding genes are regulated by a unique lncRNA in trans. In contrast, only five protein-coding genes were targeted by more than one lncRNA (Figure 4b), indicating that convergent trans regulation by multiple DE lncRNAs was very limited.

Four target genes, including plant self-incompatibility protein homologs, *JAZ5*, *PADRE*, and *SWEET4*, exhibited the most negative ndG values among the predicted trans interactions **(Figure 4c)**, indicating the strongest predicted lncRNA-mRNA binding energies. These interactions fall well beyond the primary ndG cutoff (≤-0.08) and reside in the extreme left tail of the ndG distribution, representing a small subset of energetically highly favourable RNA–RNA hybridization events. In addition to these, several other high-confidence targets with strong energetic support, including RALF-like peptides (RALFL12 and RALFL13), the exocyst subunit EXO70, cell wall-modifying enzymes (CSLC4 and XTR6), the transcription factor MYB97, and the membrane-associated protein MLO12, were identified. The ten selected trans-target genes are summarized in **Table 2**.

**Table 2.**
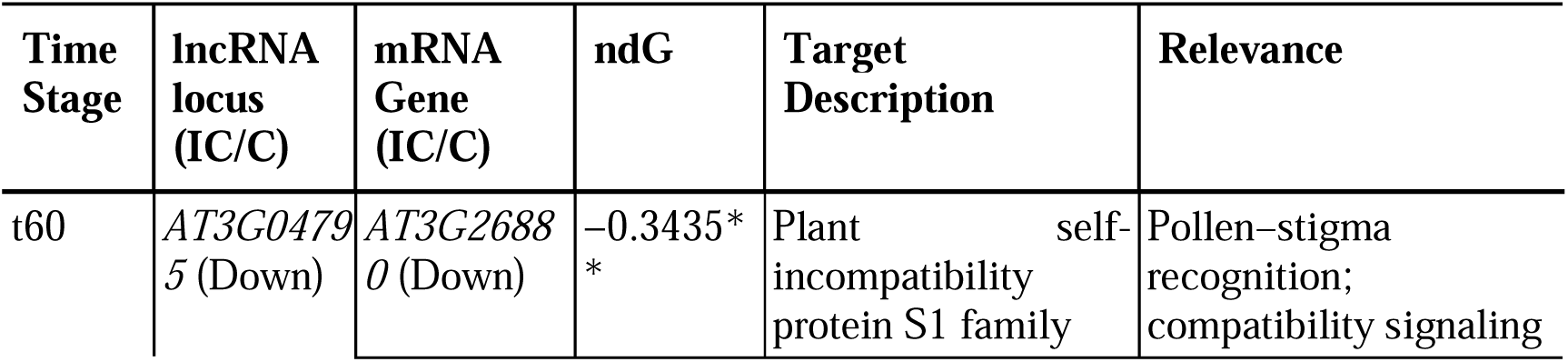
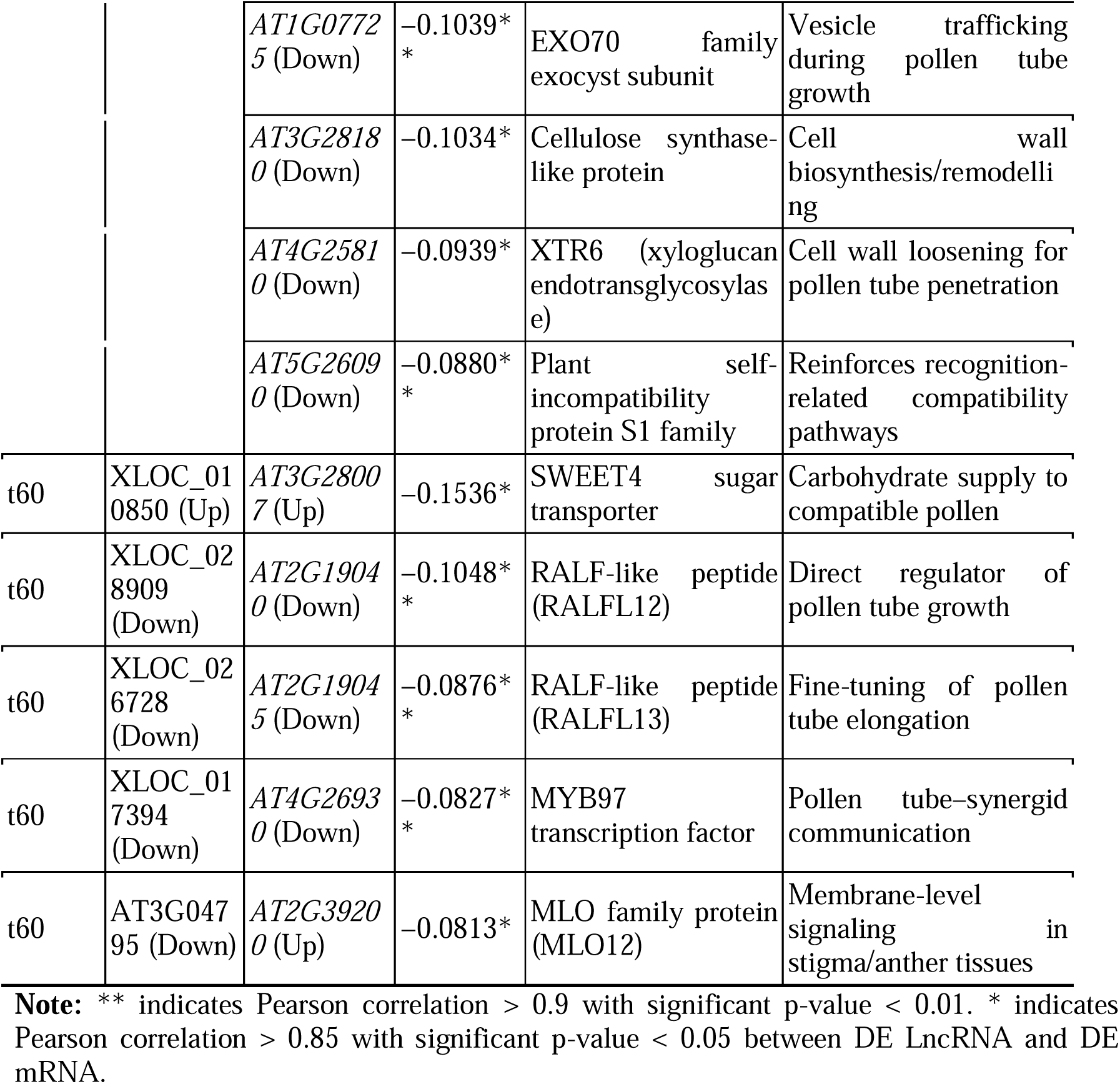
Selected trans-acting lncRNA–mRNA interaction pairs at t = 60 min (t60).

### 3.5. Late post-pollination displayed expanded network of lncRNAs, miRNAs, and mRNA targets

LncRNAs can regulate gene expression by interacting with microRNAs (miRNAs) and protein-coding genes, and lncRNA–miRNA–mRNA regulatory networks are mostly formed by competing for shared miRNAs or modulating miRNA-mediated repression (Cesana et al., 2011; Salmena et al., 2011). To investigate this regulatory mechanism during pollination responses, we constructed lncRNA–miRNA–mRNA networks by integrating DE lncRNAs, mRNAs, and known miRNAs from the miRbase database. We first mapped mRNA–miRNA interactions and then lncRNA–miRNA interactions. The analysis of the integrated lncRNA–miRNA–mRNA regulatory network identified 31 lncRNAs, 55 miRNAs, and 144 unique protein-coding genes across both time points. The early time point (10 min) showed only five lncRNAs, seven miRNAs, and 19 mRNA targets, whereas the late time point (60 min) displayed a substantially expanded network with 26 lncRNAs, 51 miRNAs, and 128 mRNA targets **(Supplementary Table S17)**.

Putative miRNA sponge interactions consist of pairs of lncRNA, miRNA, and mRNA, where lncRNA and their corresponding mRNA target show a consistent pattern of expression, either upregulation or downregulation for both, whereas opposing regulatory interactions are those where opposite expression changes between lncRNAs and their target mRNAs are found (Babaei et al., 2024). Using this criterion, we identified 28 lncRNAs that can function as potential miRNA sponges, interacting with 110 unique protein-coding genes through 49 miRNAs. This consistent pattern of expression suggests that certain lncRNAs may regulate target gene expression by sequestering shared miRNAs and thereby alleviating miRNA-mediated repression during pollination responses.

Among the identified miRNAs, ath-miR5021 and ath-miR5015b were the two most prominent miRNA hubs across both time points **(Figure 5a and 5b)**. ath-miR5021 was associated with DE 54 unique protein-coding target genes, of which 17 were upregulated and 37 were downregulated, while ath-miR5015b regulated 17 unique targets, including four upregulated and 13 downregulated genes. These hub miRNAs interacted with multiple lncRNAs and target genes, highlighting their key role in shaping lncRNA–miRNA–mRNA regulatory networks during both early and late pollination responses. The identified key miRNA hubs targeted genes involved in defense and stress responses, including *NPR3*, *PUB25*, and *TIR-NBS-LRR* immune receptors, as well as regulators of hypoxia and hormone signaling, such as *HUP17*, *Carbonic Anhydrase* (*CA2*), *JAZ5*, and *ERS2*. In addition, targets involved in pollen–pistil interaction–related processes, including *SWEET4*, *GAE1*, and β*-1,3-glucanase* (*BG5*), were also identified.

**Figure 5.**
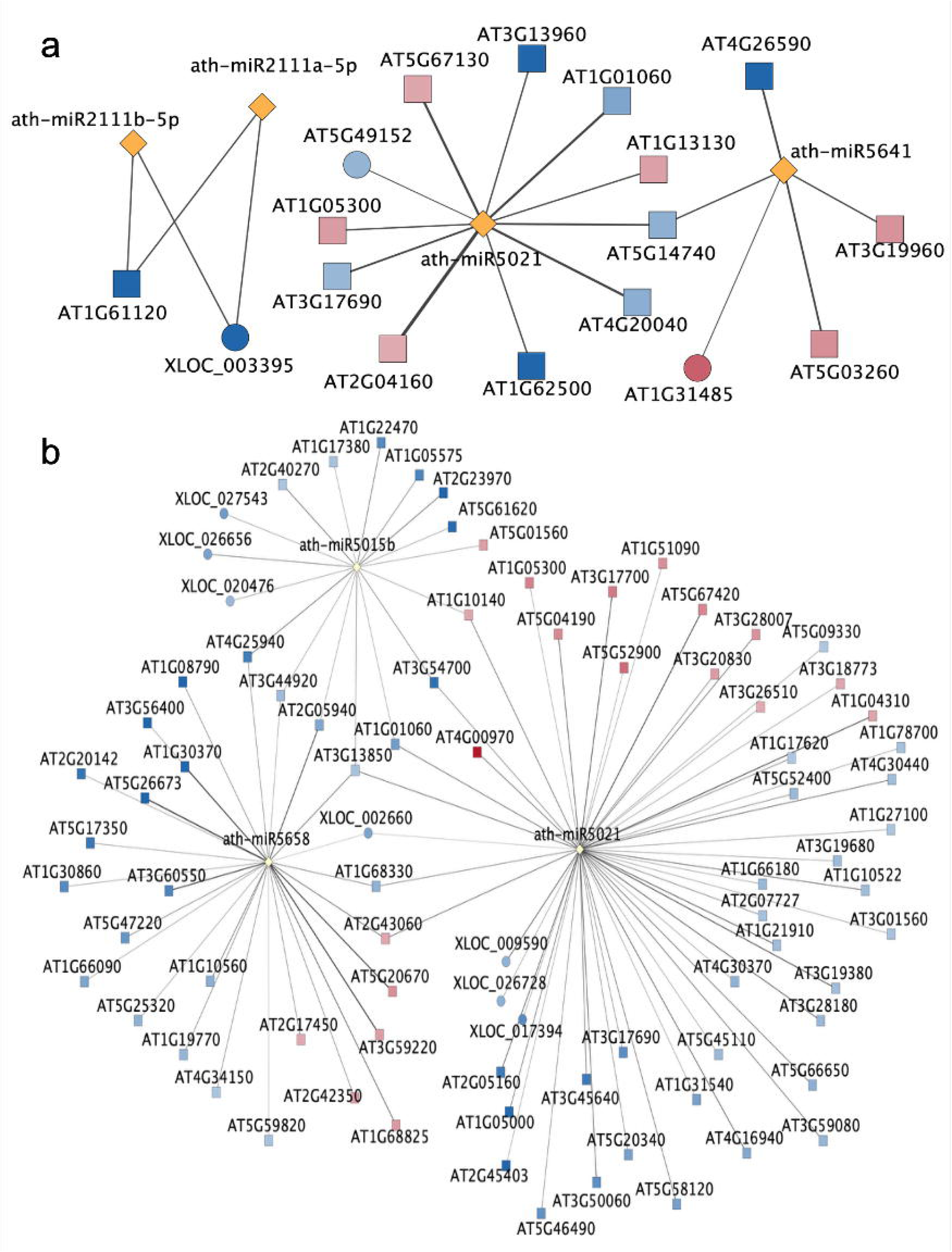
LncRNA–miRNA–mRNA interaction networks. The figure presents (a) LncRNA-miRNA-mRNA regulatory network at 10 min post-pollination. (b) LncRNA-miRNA-mRNA regulatory network at 60 min post-pollination. Only miRNAs exhibiting high-confidence interactions with multiple lncRNAs and mRNA targets were included to ensure clarity of network visualization. Circles represent lncRNAs, rectangles represent mRNA targets, and yellow diamonds represent miRNAs. Node colour indicates relative expression changes between incompatible and compatible pollination (IC/C), with increasing blue intensity denoting down-regulation and increasing red intensity denoting up-regulation.

### 3.6. Heatmap displayed progressive increase in target gene expression from early to late post-pollination

Post-pollination treatments at 60 min showed higher numbers of expressed lncRNAs, DE lncRNAs, DE mRNAs, and associated target genes than at 10 min. The analysis of cis, trans, and lncRNA–miRNA–mediated interactions showed a notable increase in the number of differentially regulated target genes at 60 min compared to 10 min (**Figure 6a**). Gene Ontology (GO) enrichment analysis was performed using target genes from interaction pairs between DE lncRNAs and DE mRNAs. The enriched GO terms included hypoxia response, stress regulation, defense mechanisms, and hormone signaling **(Figure 6b**, **Supplementary Table S18)**.

**Figure 6.**
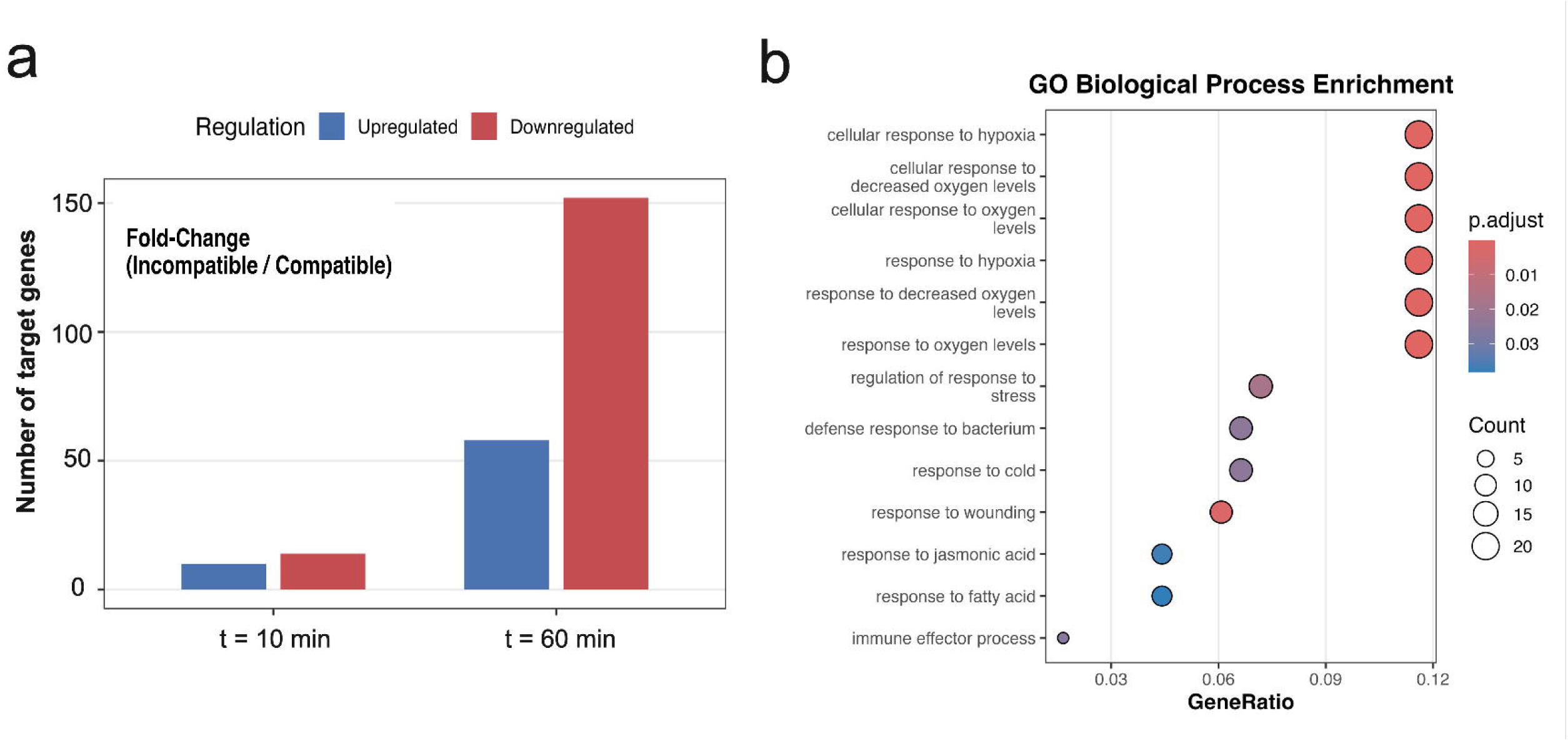
Temporal dynamics and functional enrichment of differentially regulated target genes during compatible and incompatible pollination of *Arabidopsis thaliana*. The figure shows (a) The number of upregulated and downregulated differentially expressed (DE) target genes at 10 and 60 min post-pollination, based on fold change between incompatible and compatible conditions (IC/C). (b) Gene Ontology (GO) biological process enrichment analysis of the target genes, visualized as a dot plot. Dot size represents the number of genes associated with each GO term, while colour intensity indicates the adjusted p-value. The x-axis denotes the gene ratio for each enriched biological process.

The global expression changes in lncRNA-associated target genes were visualized using a heatmap across both time points and pollination conditions **(Figure 7),** which included targets identified through cis, trans, and miRNA-mediated regulatory interactions across 12 samples. The target genes were categorized into four distinct groups based on the time point at which they were identified and their expression levels under pollination conditions. These categories included targets upregulated at 10 minutes (t10 Up), downregulated at 10 minutes (t10 Down), upregulated at 60 minutes (t60 Up), and downregulated at 60 minutes (t60 Down). **(Supplementary Table S19)**. The targets identified at 60 min showed more distinct and coordinated expression changes, particularly under incompatible conditions, consistent with the increased number of differentially regulated targets observed at 60 min (**Figure 7**). Together, this heatmap highlights the temporal specificity and progressive reprogramming of target gene expression during the transition from early to late pollination responses.

**Figure 7.**
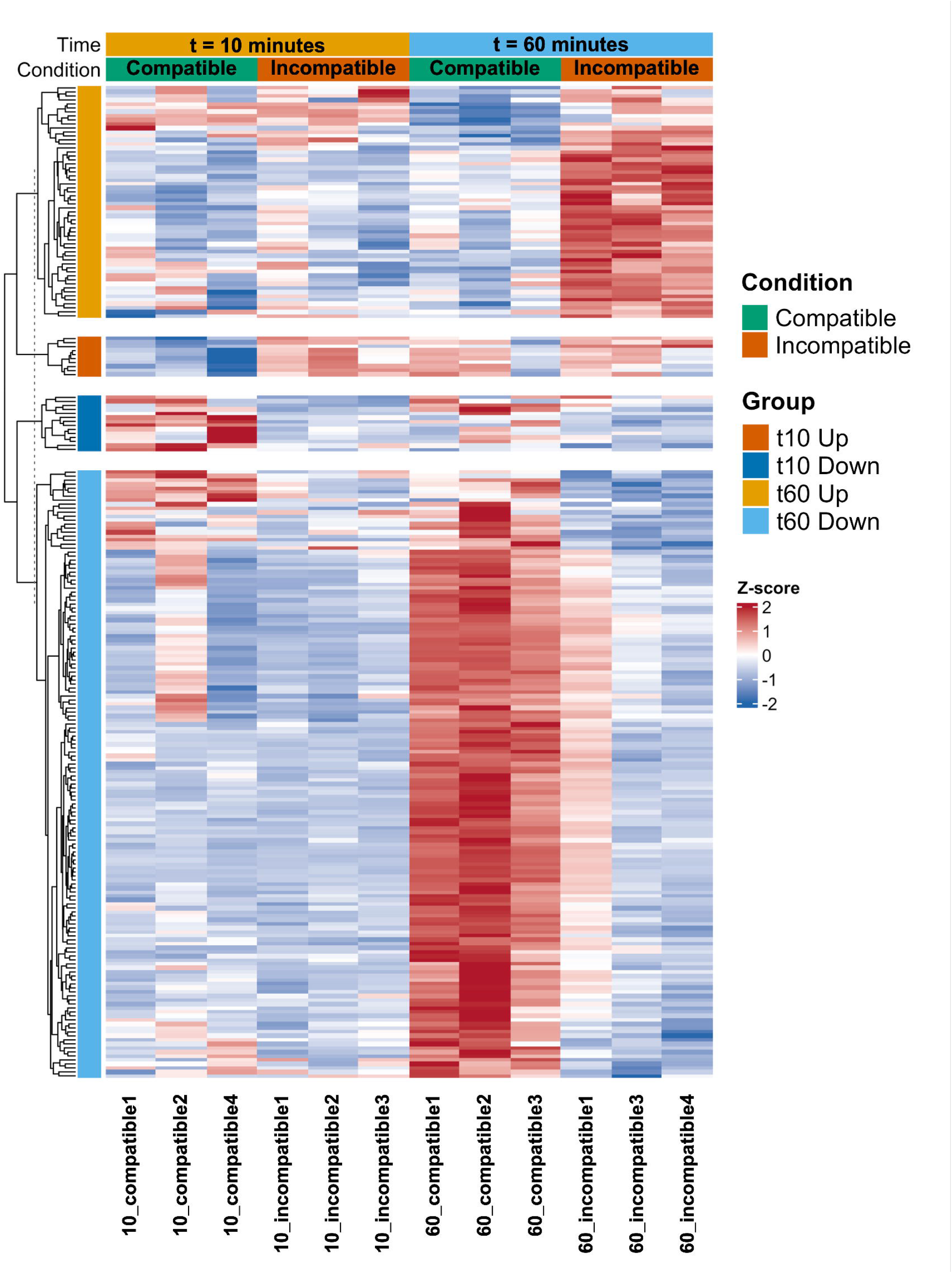
Global expression patterns of target genes across time points and pollination conditions. Heatmap showing gene-wise Z-score scaled variance-stabilized expression of all predicted target genes derived from cis-, trans-, and miRNA-mediated regulatory interactions. The heatmap includes all 12 samples representing compatible and incompatible pollination conditions at 10 and 60 min post-pollination. Genes are grouped into four sections based on their direction of regulation and the time point at which each target was identified: targets upregulated at 60 min (t60 Up), targets upregulated at 10 min (t10 Up), targets downregulated at 10 min (t10 Down), and targets downregulated at 60 min (t60 Down), as determined by fold change between incompatible and compatible conditions (IC/C). Column annotations indicate time point and pollination condition, while colours represent relative expression levels, with red indicating higher and blue indicating lower expression relative to the gene-wise mean.

## 4. Discussion

During self-incompatibility responses in *Brassicaceae*, a cascade of signaling events is triggered. The stigma localized S-locus receptor kinase (SRK) recognizes the S-locus cysteine-rich protein (SCR/SP11) from self pollen, activating the downstream E3 ubiquitin ligase ARC1, leading to degradation of compatibility factors (Abhinandan et al., 2022; Abhinandan et al., 2023; Bhalla et al., 2025a). In parallel, self-pollination induces ROS production through activation of FERONIA-mediated signaling, which disrupts pollen hydration and tube growth, resulting in rejection of self-pollen. In contrast, during compatible pollination, pollen coat proteins (PCP-Bs) suppress ROS production by competing with RALF23/33 peptides for interaction with the ANJEA-FERONIA receptor complex, thereby promoting pollen acceptance (Abhinandan et al., 2022; Bhalla et al., 2025b). To investigate the regulatory role of lncRNA in these pathways and identify novel molecular interactors, we performed an integrative analysis of lncRNAs, their mRNA targets, and associated miRNA networks using RNA-seq data. The datasets used in this study consisted of three replicates each for compatible and incompatible pollination at two time points, 10 min and 60 min post-pollination (Kodera et al., 2021). Since the stigmatic tissues were pollinated with both compatible and incompatible pollen, the RNA-seq data capture transcriptional responses from the stigma and from the pollen as it grows through the stigmatic tissue. The replicate that diverged from the three replicates identified by principal component analysis (PCA) was excluded from further analysis to maintain the integrity and robustness of our analysis. A stringent three-step filtering approach using CPC2, CPAT, and LncFinder identified 9888 lncRNA candidates, including 1,533 novel lncRNA transcripts spanning 1,073 lncRNA loci. This approach minimized false positives and enhanced dataset reliability.

The predominance of sense-genic lncRNAs suggests their potential involvement in regulating nearby protein-coding genes associated with pollination responses. LncRNAs were unevenly distributed across chromosomes; likely due to evolutionary selection pressures or to their functional specialization. However, the unique properties of lncRNAs may arise due to their distinct roles within the transcriptomic landscape. These findings highlight lncRNA diversity in reproductive processes and provide a foundation for future investigations into their regulatory mechanisms in gene expression. Their distinct expression patterns between compatible and incompatible pollination further support the notion that lncRNAs act as active regulators rather than byproducts of transcription.

Differential expression analysis of lncRNA loci revealed that at 10 min, only 11 loci were expressed, whereas at 60 min, 31 loci were expressed, among which the majority of differentially expressed lncRNAs were downregulated, with the highest reduction of 7.8-fold (**Figure 3c**). These results suggest that later stages of pollination are associated with the activation of additional lncRNAs, which may have a significant regulatory role in reproductive success. The presence of a higher number of expressed lncRNA transcripts under compatible pollination at both t = 10 min and t = 60 min time points of post-pollination suggests a potential role for these lncRNAs in promoting compatibility and inhibiting responses that might lead to pollen rejection. A similar trend of downregulation was also observed in differentially expressed mRNAs at similar time points, with the majority downregulated. The downregulation of these lncRNAs may reflect their role in fine-tuning gene expression, while the similar pattern for mRNA levels may help optimize compatibility. The analysis of significant cis-acting lncRNA–mRNA interaction pairs at t = 10 min and t = 60 min of post-pollination showed that, at both time points, the lncRNA XLOC_004982 exhibited a strong positive correlation with the mRNA *AT1G14870* (*PCR2*), both of which were downregulated. *PCR2* belongs to the Arabidopsis PCRs, a small gene family of 12 members, including PCR11, which is expressed in pollen and belongs to the same clade as PCR2 (Song et al., 2010). PCR2 encodes a protein involved in zinc transport and detoxification, suggesting a potential regulatory role for XLOC_004982 in metal ion homeostasis during the initial stages of pollen tube growth. It would be interesting to determine whether PCR2 is directly or indirectly involved in plant reproductive processes, especially during pollination.

The analysis of differentially expressed genes (DEGs) in the pistil transcriptomes of *Arabidopsis thaliana* and *Arabidopsis halleri* during self-pollination, interspecific pollination, and pathogen infection with *Fusarium graminearum* showed that up to 79% of down-regulated genes were shared between pollination and pathogen response (Mondragón-Palomino et al., 2017). Meanwhile, interspecific pollination of *A. thaliana* upregulates thionins and defensins, which are involved in the defense mechanism. Consistent with this, our analysis identified several stress-and defense-related genes among lncRNA targets, including members of the MEKK subfamily (*AT1G07128*), which are implicated in wound-induced signaling, indicating a connection between physical stress responses and pollination. Downregulation of genes such as Chalcone-flavanone isomerase (AT3G51238) and DREB subfamily member A-5 (AT1G21910) proteins suggests a strategic shift toward prioritizing reproductive processes over stress responses. The negative correlation between XLOC_029258 and AT5G40020 at the early time point suggests a complex relationship in which the lncRNA may influence the expression of a pathogenesis-related protein. Similarly, the negative correlation between AT1G07128 and AT1G07150, and the strong correlation between XLOC_029189 and AT5G39050 at 60 min, indicate that the regulatory mechanism involved defense responses. The interaction between pollination and pathogen response indicates the complex relationship between plant reproductive strategies and defense mechanisms. This emphasizes the role of lncRNAs as key regulators of gene expression through significant lncRNA–mRNA interactions.

Trans-acting lncRNA-mRNA interaction analysis identified key players involved in pollination. All these targets were detected at 60 min post-pollination. The members of the exocyst complex regulate polarized secretion and are important for the acceptance of compatible pollen in both Brassica and Arabidopsis (Samuel et al., 2009; Safavian et al., 2015). Among these members, EXO70A1, EXO70A2 and EXO70C2 are essential for pollen maturation, germination, and tube growth (Samuel et al., 2009; Marković et al., 2020; Saccomanno et al., 2021). Our results identified the interaction of lncRNA AT3G04795 and EXO70 family exocyst subunit (*AT1G07725*), suggesting the role of these interactions in vesicle trafficking required pollen tube growth. The downregulation of the lncRNA AT3G04795 is linked to several mRNA targets, including *AT3G26880* and *AT5G26090*, which encode a self-incompatibility protein S1 (SPH family member). The SPH family was initially identified in the self-incompatibility response of the field poppy (Rajasekar et al., 2019). *AT3G26880* and *AT5G26090* are highly expressed in the pollen tube of Col-0 (https://evorepro.sbs.ntu.edu.sg/), suggesting that lncRNA AT3G04795 negatively regulates the expression of these genes, disrupting the pollen tube growth during incompatible pollination. The lncRNA XLOC_010850 interacts with *AT3G28007* (*SWEET4*). The SWEET4 homolog in Arabidopsis, SWEET5, functions in later stages of pollen development, whereas SWEET4 is involved in sugar transport to axial tissues during plant growth and development (Liu et al., 2016). This interaction suggests that lncRNA controls nutrient allocation during pollen development. RALF family members (e.g., RALF4/9) and their pollen-tube receptors Buddha’s Paper Seal 1 and 2 (BUPS1/2) are essential for normal pollen tube growth (Ge et al., 2017). *RALFL12* and *RALFL13* displayed higher expression in Col-0 pollen tubes (https://evorepro.sbs.ntu.edu.sg/). In our study, XLOC_028909 and XLOC_026728 (both downregulated) target *RALFL12/13,* suggesting their role during compatible pollination.

The MLO family proteins (MLO1, 5, 9, and 15) act as Ca^2+^ channels, enabling influx to sustain pollen tube integrity. RALF peptides bind pollen-tube receptors to activate these channels and establish a Ca^2+^ gradient (Gao et al., 2023). Our study shows that downregulation of lncRNA (AT3G04795) coincides with upregulation of target *MLO12,* implying negative regulation. Previous studies have reported a shared pathway for the pathogen response and pollination (Kessler et al., 2010; Mondragón-Palomino et al., 2017). Mutations in MLO gene family members, especially MLO2, MLO6, and MLO12, limit powdery mildew colonization and influence interactions with a range of other phytopathogens. In our study, the interaction between lncRNA AT3G04795 and MLO12 supports an integrated regulatory framework connecting pollination and disease resistance, with MLO12 serving as a key component in this defense pathway and possibly playing a role in maintaining pollen tube integrity, as do other MLOs.

MYB transcription factors regulate male reproductive development in flowering plants. In Arabidopsis, loss of pollen-specific MYB97, MYB101, and MYB120 reduces the expression of pollen-tube-expressed genes and disrupts pollen ability to burst upon reaching a synergid cell, ultimately impairing fertilization (Leydon et al., 2014). These MYB genes are localized to the pollen tube nucleus and are involved in coordinating pollen tube gene expression during pistil growth. In our study, lncRNA XLOC_017394 targets *MYB97* (*AT4G26930*), and both transcripts are downregulated, suggesting that lncRNA XLOC_017394 may be a key regulator in modulating *MYB97* expression, ultimately affecting pollen tube development and male reproductive success during the early pollen-pistil interaction.

Among the predicted trans interactions, four target genes, including plant self-incompatibility protein homologs *JAZ5*, *PADRE*, and *SWEET4,* exhibited the most negative ndG values **(Figure 4c)**. The jasmonate hormone (JA) plays a critical role in both plant defense and reproductive development. In Arabidopsis, plants deficient in JA-biosynthesis or signaling are male-sterile, displaying defects in stamen and pollen development. In carrot protoplasts, the bHLH transcription factor MYC5 activates GUS reporter genes driven by the JAZ5 promoter, and MYC5 likely acts together with other transcription factors to induce *MYB21* and other players required for male fertility (Figueroa et al., 2015).

LncRNAs are long, diverse regulatory RNAs transcribed by RNA polymerase II and are processed like mRNAs (capping, splicing, and polyadenylation). In contrast, miRNAs are short (∼21 nt) RNAs derived from primary transcripts (pri-miRNAs) that are processed in plants by Dicer-like 1 (DCL1) into mature miRNAs, which are then introduced into the RNA-induced silencing complex (RISC) to guide sequence-specific binding to target mRNAs, resulting in mRNA cleavage (Yu et al., 2026). In our study, ath-miR5021 and ath-miR5015b were identified as the two most prominent miRNA hubs across both time points **(Figure 5a and 5b),** targeting several genes, including those involved in differentiation and plant reproductive processes. These miRNA hubs also regulate several genes with established roles in defense and stress-associated pathways. Among the targets is *NPR3*, a salicylic acid (SA) receptor that binds SA with different affinities and functions as an adaptor for the Cullin 3 ubiquitin E3 ligase to mediate SA-regulated degradation of related receptor NPR1, thereby contributing to systemic acquired resistance in plants (Fu et al., 2012). Additional targets include the members of the Plant U-box E3 ligase gene family, such as PUB25, which confers freezing tolerance in plants and is highly expressed in root and internode tissues (Wang et al., 2019; Karthik et al., 2025). Other targets also included the defense and immune receptor genes, including *JAZ5* and *TIR-NBS-LRR* immune receptor genes. These hubs also target genes such as β*-1,3-glucanase* (*BG5*) and *GAE1*. In Arabidopsis, a single somatic ovule differentiates into the female germline, which is protected by a water-insoluble polysaccharide, β−1,3-glucan, which restricts pathogen invasion while also promoting pollen development. Thus, germline β−1,3-glucan insulates the primary germline cell and contributes to the success of downstream female gametogenesis. Other targets, such as *Glucuronate 4-epimerases* (*GAE*s), like *GAE1*, convert UDP-D-glucuronic acid to UDP-D-galacturonic acid, catalyzing a key step in pectin biosynthesis, and are essential for maintaining the structure and integrity of plant cell walls. GAE1 also contributes to pathogen resistance, as *GAE1* and *GAE6* mutants exhibit compromised disease resistance (Bethke et al., 2016).

Gene Ontology (GO) analysis revealed significant enrichment in biological processes related to adaptive and immune responses during stress and pathogen attack. Enriched GO terms included GO:0071456 (cellular response to hypoxia), GO:0036294 (cellular response to decreased oxygen levels), and GO:0009611 (response to wounding), suggesting the involvement of lncRNAs and their network in adaptive responses. Other GO terms, such as GO:0002252 (immune effector process) and GO:0042742 (defense response to bacterium), which are associated with immune responses, were also enriched. Overall, the mRNA targets of lncRNA and miRNA hubs, together with GO term analysis, reveal a gene interaction network primarily involved in stress and defense responses, consistent with previous studies (Mondragón-Palomino et al., 2017; Kodera et al., 2021). However, novel lncRNAs with unknown functions in plant reproduction, especially during compatible and incompatible pollination, require further investigation and functional validation.

## 5. Conclusion

In this study, we identified a high-confidence set of both novel and annotated lncRNAs expressed during compatible and incompatible pollination in *Arabidopsis thaliana* at early and late post-pollination stages. Although comparable numbers of lncRNAs were detected at both time points, incompatible pollination induced a strong, time-dependent shift in lncRNA regulation, characterized by predominantly downregulated and largely stage-specific differentially expressed lncRNAs at the later stage. Cis-regulatory associations of lncRNAs were relatively limited in number but exhibited strong expression concordance with neighboring protein-coding genes, suggesting tightly coordinated local regulation. In contrast, the trans regulatory interactions were markedly enriched at the late stage and were driven by a small number of highly connected lncRNAs targeting genes involved in self-incompatibility, stress, and hormone-related pathways. Integration of miRNA-mediated regulation revealed an additional regulatory layer, with expanded lncRNA-miRNA-mRNA networks at a later stage and a predominance of putative miRNA-sponge interactions. Analysis of lncRNA-associated target genes showed a strong temporal specificity and coordinated transcriptional changes in global expression analysis, which shows a transition from early signaling events to broader regulatory responses during incompatible pollination.

## Statements

### Data availability statement

The datasets used in this study and the newly generated datasets are provided in Supplementary Tables S1 to S19.

### Author contributions

NP: Formal analysis, Methodology, Writing – original draft, NDG: Writing – review & editing, SS: Conceptualization, Supervision, Writing – review & editing.

### Funding

This project was supported by the Ramalingaswami Re-entry Fellowship, a grant from ANRF (ARG/2025/000937/LS), and by a start-up grant from the Indian Institute of Technology Gandhinagar to SS.

## Acknowledgments

We are grateful to Dr. Isabelle Fobis-Loisy, INRAE, France, for the raw RNA-seq data used in this study and useful suggestions. We are thankful to the Indian Institute of Technology Gandhinagar for a post-doctoral fellowship to NG. We also acknowledge DBT for the Ramalingaswami Re-entry fellowship grant, SERB, ANRF, and the Indian Institute of Technology Gandhinagar for a start-up grant to SS.

## Conflict of interest

The authors declare no conflict of interest

## Notes

### Competing Interest Statement

The authors have declared no competing interest.

