## Supplementary Figure 1 and 2 for "Characterization of long non-coding RNAs during compatible and incompatible pollination in *Arabidopsis thaliana*"

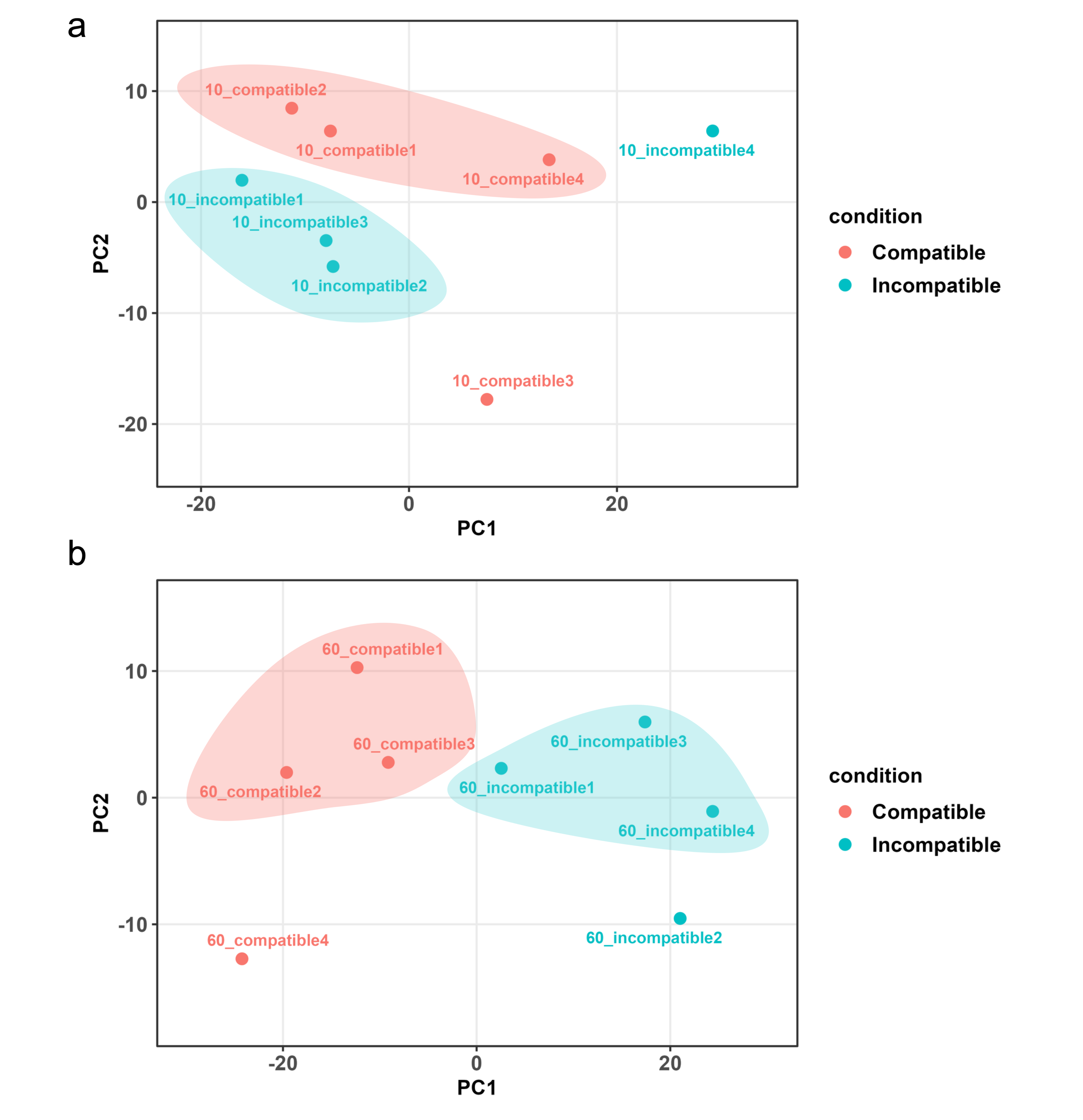

**Supplementary Figure S1. Principal component analysis (PCA) of RNA-seq samples at early and late pollination time points.** (a) PCA of variance-stabilized RNA-seq expression data at t = 10 min after pollination and (b) PCA at t= 60 min after pollination. PCA was performed on variance-stabilized counts generated using the DESeq2 pipeline. Each point represents an individual biological replicate, colored by pollination condition (Compatible and Incompatible). The percentage of variance explained by PC1 and PC2 is indicated on the axes. One outlier replicate per condition was removed prior to PCA visualization: 10_incompatible2 and 10_compatible3 at 10 min, and 60_incompatible2 and 60_compatible4 at 60 min. PCA was used to assess sample relatedness and consistency among biological replicates within and between pollination conditions at each time point.

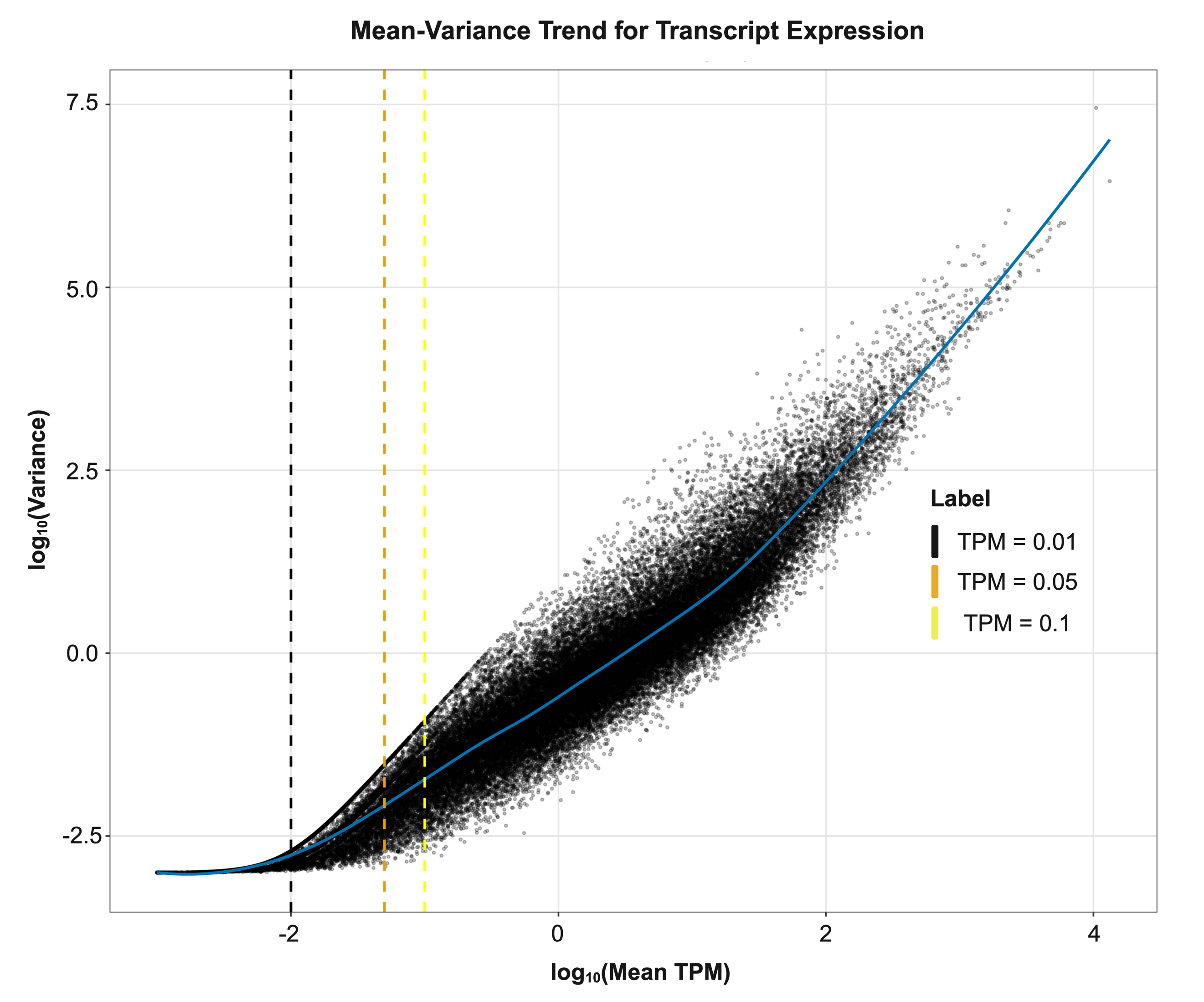

**Supplementary Figure S2.** **Mean–variance relationship of transcript expression at early and late pollination time points.** Mean–variance trend analysis of RNA-seq expression data of Compatible and Incompatible pollination. Each point represents a transcript, plotted as log₁₀ (mean TPM) versus log₁₀ (variance) across biological replicates. The blue curve indicates the locally weighted regression (LOWESS) fit capturing the global mean–variance relationship. Vertical dashed lines denote commonly used TPM expression thresholds: TPM = 0.01 (black), TPM = 0.05 (orange), and TPM = 0.1 (yellow). Transcripts with very low mean expression (TPM ≤ 0.05) exhibit elevated variance relative to their mean, consistent with increased variability at low expression levels. As mean expression increases, the mean–variance relationship becomes progressively smoother, with improved coupling observed around TPM > (0.05 – 0.1). This analysis was used to inform the choice of expression thresholds for downstream analysis.
